# Histotripsy-induced vascular remodeling in the tumor microenvironment enhances local and off-target intratumoral drug delivery

**DOI:** 10.64898/2026.09.18.752667

**Authors:** Heineken Queen, Brian Song, Hanna Kim, Reliza McGinnis, Jiayu Liu, Aichata Sanogo, Katherine Buglak, Chaitanya Karanam, Allison Bontrager, Tejaswi Worlikar, Zhen Xu, Anutosh Ganguly, Clifford S. Cho

## Abstract

Solid tumors are known to be refractory to treatment due to a poorly perfused microenvironment that blocks immune cell infiltration and effective drug delivery. Histotripsy is a non-invasive focused ultrasound modality that generates cavitation microbubbles to mechanically disrupt solid tumors, resulting in decreased hypoxia and stimulation of tumor-targeted adaptive immune responses. In this study, we aim to investigate whether post-histotripsy hypoxia reduction and cytotoxic immune cell infiltration are associated with normalization of tumor vasculature and enhanced perfusion. Structural analysis of both treated and contralateral off-target tumors following partial histotripsy revealed significant vascular remodeling, enhanced vascular integrity, and decreased vascular leakage indicative of vascular normalization in multiple *in vivo* models of melanoma, hepatocellular carcinoma, and pancreatic adenocarcinoma. These effects appeared antigen-specific, as no remodeling was observed in contralateral off-target tumors of discordant pathology. Mechanistically, CXCR3+CD8+ T cells played a role in mediating these vascular changes, as genetic knockout of CD8 and pharmacological antagonism of CXCR3 impaired both remodeling and normalization. Additionally, significant changes in endothelial CXCR4 and Angiopoietin-1 and -2 expressions were observed following histotripsy, indicating their potential involvement in promoting vascular normalization and remodeling. Significant vascular remodeling was accompanied by enhanced tumor perfusion and intratumoral delivery of chemotherapeutic agents and therapeutic monoclonal antibodies. These findings suggest that histotripsy not only creates a more favorable microenvironment for immunotherapy, but also optimizes drug delivery into solid tumors. Combinatorial approaches using histotripsy with chemotherapy or immunotherapy may be a novel therapeutic strategy to overcome current limitations of conventional solid tumor therapies.

**Summary:** Histotripsy reorganizes the tumor vasculature of solid cancers, promoting enhanced drug delivery into treated and non-targeted distant tumors.

## INTRODUCTION

The highly hypoxic state of the tumor microenvironment (TME) gives rise to the aberrant vasculature of solid tumors, substantially impeding immune infiltration and drug delivery and contributing to therapeutic resistance and tumor progression (*1–3*). Tumor cells respond to chronic hypoxia with transcriptional programs that enable them to thrive under the harsh conditions of the TME, including upregulated expression of vascular endothelial growth factor (VEGF) that promotes the formation of new blood vessels from pre-existing ones (known as angiogenesis) (*4–6*). Several mechanisms regulate tumor angiogenesis: the CXCR4:CXCL12 signaling axis indirectly promotes blood vessel sprouting by inducing expression of VEGF (*7–10*); angiogenesis is also initiated by a switch from angiopoietin-1 (Angpt1) expression (which drives endothelial cell maturation, migration, and survival) to angiopoietin-2 (Angpt2) expression (which antagonizes Angpt1 binding to the Tie-2 receptor and causes vessel destabilization and regression) (*11–13*).

Rapid angiogenesis results in the formation of immature and dysfunctional blood vessels characterized by high vessel density, disrupted endothelial junctions, lack of pericyte coverage, and anatomic malformations that lead to increased interstitial fluid pressure and poor perfusion (*14–17*). Persistent structural and functional vessel abnormalities perpetuate a vicious cycle of chronic hypoxia and aberrant angiogenesis, supporting tumor growth, metastasis, and treatment failure (*18–19*). Thus, one appealing strategy to treat solid tumors is to target the tumor vasculature. Indeed, efforts have been made to induce so-called vascular normalization, re-organizing and stabilizing dysfunctional vasculature in the hopes of improving blood flow, drug penetration, and immune cell infiltration (*20–22*). Antiangiogenic drugs including bevacizumab, sunitinib, and axitinib are currently being used in the clinic to starve cancer cells by blocking VEGF-mediated tumor angiogenesis; however, therapy resistance often occurs via redundancies in angiogenic signaling pathways, recruitment of bone marrow-derived endothelial progenitor cells, and adaptation to hypoxia caused by vessel pruning (*23–27*).

Histotripsy is a high-precision, non-invasive, and non-thermal focused ultrasound ablation technique that recently received FDA approval for the treatment of primary and metastatic liver tumors (*28–29*). In our previous study, we demonstrated that histotripsy abrogates hypoxia response elements (HREs) including hypoxia-inducible factor-1 alpha (HIF-1α) and VEGF to stimulate tumor-directed immune responses (*30*). The present study examines whether the hypoxic and immune effects of histotripsy are linked to normalization of tumor vasculature. Here, we aim to determine whether histotripsy can reorganize the tumor vasculature to potentiate the efficacy of immunotherapy and chemotherapy by improving drug delivery.

## RESULTS

### Histotripsy inhibits tumor growth and alters intratumoral vascular architecture

We sought to investigate how the vasculature of solid tumors is affected by histotripsy in treated and in contralateral, off-target tumors. C57BL/6 mice bearing bilateral flank B16F10, Hepa1-6, or mT4 tumors were treated with sham or subtotal 70% histotripsy tumor treatment when tumors reached 8-10 mm in maximal diameter (about 7 days after tumor inoculation). As shown in **Supplemental Fig. S1**, partial histotripsy decreased the tumor growth kinetics for all tumor types, at both treated and contralateral off-target tumors. A morphometric assessment of tumor vasculature was conducted one week after histotripsy for B16F10 and Hepa1-6 tumors, and ten days post-treatment for mT4 tumors (**Fig. 1A–C**). Angiogenesis indicators including the number of extremities (branch points) and the total length of isolated branches were reduced by half in treated B16F10 and Hepa1-6 tumors compared to controls; notably, similar reductions were also observed in contralateral, off-target tumors after histotripsy (**Fig. 1D–F**). While morphometric features of angiogenesis did not change significantly in mT4 tumors, a substantial reduction in vessel density was observed in all tumor models (**Fig. S2A**; **Fig. 1G–I**; **Table 1**). To determine if histotripsy leads to morphometric changes in a more clinically relevant setting, we established an orthotopic model of hepatocellular carcinoma (HCC) by implanting McA-RH7777 cells into the liver of immunocompetent Sprague-Dawley rats. Morphometry analysis of tumor vasculature revealed significant changes in all morphometric features, including a significant reduction in the number of extremities, number of isolated segments, total isolated branches length, and total length after histotripsy (**Fig. S2B–C**). Together, these observations demonstrate that histotripsy results in significant alterations in angiogenesis-associated features of vascular density and organization in both targeted and distant, off-target tumors.

**Figure 1.**
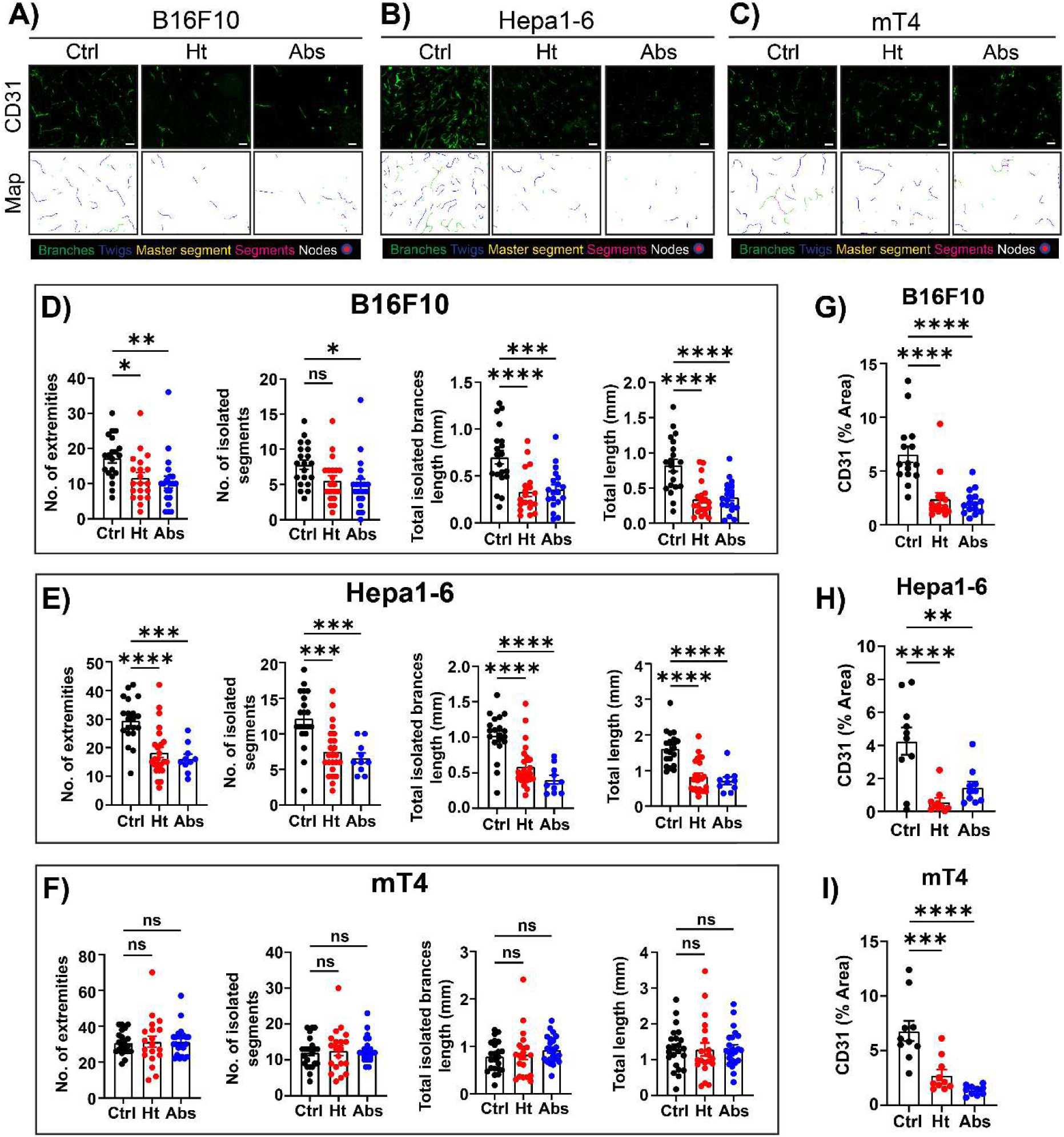
Histotripsy alters the vascular architecture of different tumor-types. (**A–C**) Tumor endothelium staining of murine melanoma (B16F10), hepatocellular carcinoma (Hepa1-6), and pancreatic adenocarcinoma (mT4), respectively, with maps showing different features of angiogenesis: branches, twigs, master segment, segments, and nodes. Scale bars represent 50µm. (**D–F**) Quantification of angiogenic features in control (Ctrl), histotripsy-treated (Ht), and non-treated contralateral (Abs) tumors. (**G–I**) Vascular density of control (Ctrl), histotripsy-treated (Ht), and contralateral (Abs) tumors as measured by percentage of CD31 (ns = not significant; *p < 0.05; **p < 0.01; ***p < 0.001; ****p < 0.0001; error bars represent standard error of the mean (SEM)).

To investigate how unilateral histotripsy affects tumor vascular integrity, immunofluorescence staining was conducted on both targeted and contralateral off-target tumors from bilateral B16F10, Hepa1-6, and mT4 models using CD31 as an endothelial marker, apelin to evaluate pericyte recruitment, and desmin for vessel maturity and stability. Analysis of apelin showed a significant increase in pericyte coverage in both targeted and off-target tumors after histotripsy treatment (**Fig. 2A–D**). Differences in the extent of pericyte coverage between B16F10 and Hepa1-6 models were noted, but these remained statistically significant in both models (**Table 2**). The proportion of desmin-positive tumor vessels rose markedly across all tumor types after histotripsy, except in contralateral, off-target Hepa1-6 and mT4 tumors (**Fig. 2E–G**, **K–M**; **Table 3**). To quantitate blood vessel leakage, fibrin deposition surrounding the tumor vasculature was analyzed in both targeted and contralateral off-target tumors. Image analysis revealed a significant reduction in fibrin deposition in targeted and off-target Hepa1-6 and mT4 tumors after histotripsy (**Fig. 2H–J**, **N–P**; **Table 3**). As shown in **Fig. 2N**, baseline fibrin leakage in B16F10 tumors was already low prior to histotripsy; while post-histotripsy fibrin deposition did not significantly decrease, a downward trend was observed. **Table 4** shows the summary of vascular remodeling in histotripsy-treated and off-target contralateral tumors. Notably, the local and abscopal vascular anatomic and physiologic effects of histotripsy appeared strongest in Hepa1-6 tumors.

**Figure 2.**
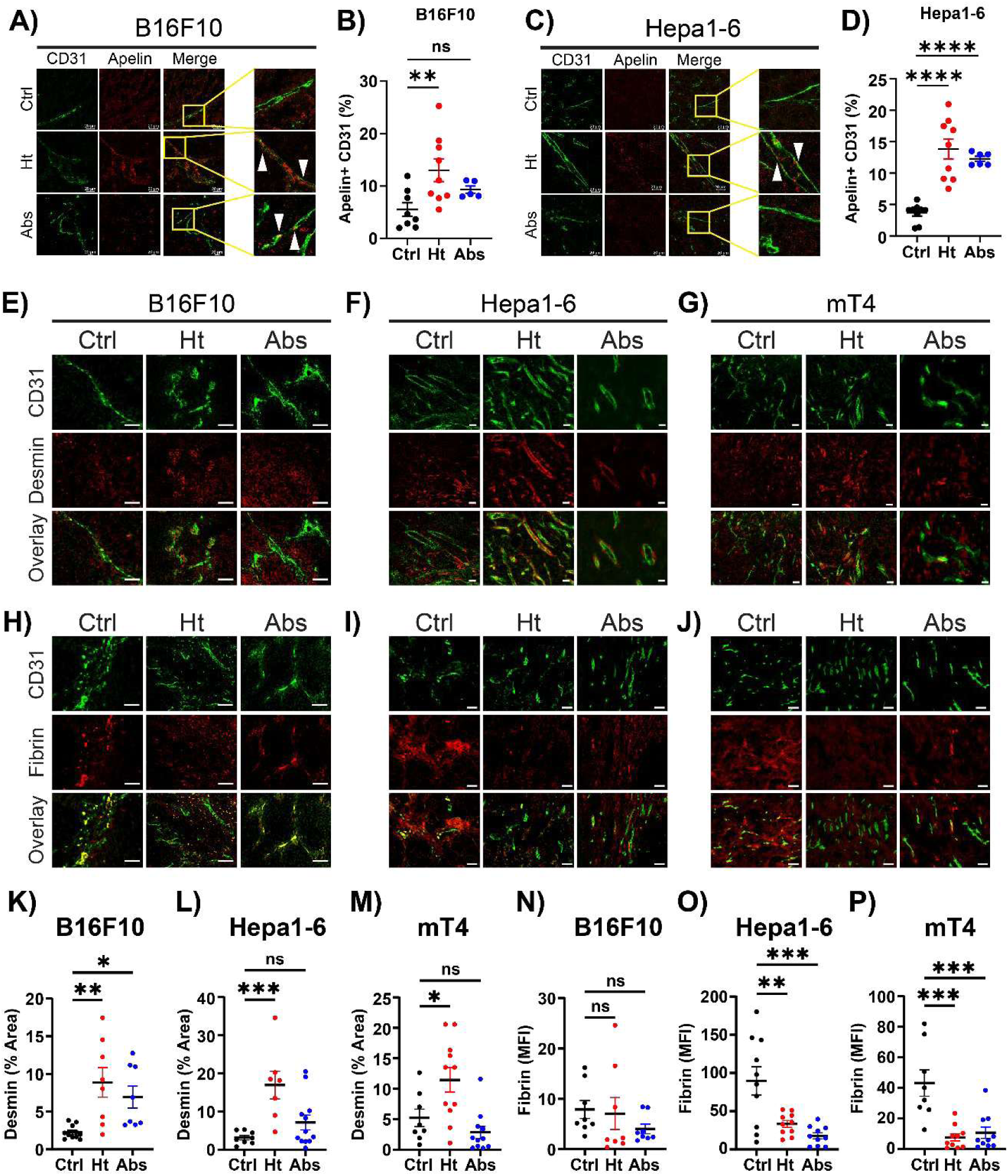
Histotripsy induces vascular normalization in solid tumors. (**A**) Immunofluorescence (IF) staining of control, histotripsy-treated, and non-treated contralateral B16F10 tumors with tumor endothelium (CD31) and pericyte marker (Apelin). (**B**) Quantification of pericyte coverage of the tumor endothelium from (**A**). (**C**) IF staining of Hepa1-6 tumors with tumor endothelium (CD31) and pericyte marker (Apelin). (**D**) Quantification of pericyte coverage of tumor endothelium from (**C**). (**E–G**) Desmin staining of control (Ctrl), treated (Ht), and non-treated contralateral (Abs) melanoma, hepatocellular carcinoma, and PDAC tumors, respectively, to indicate mature vasculature. (**H–J**) Fibrin staining of control and treated tumors to indicate vasculature leakiness. Scale bars represent 50µm unless stated otherwise. (**K–M**) Quantification of Desmin from (**E–G**). (**N–P**) Quantification of Fibrin from (**H–J**) using mean fluorescence intensity (MFI) (ns = not significant; *p < 0.05; **p < 0.01; ***p < 0.001; ****p < 0.0001; error bars represent standard error of the mean (SEM)).

Given that histotripsy seemed to improve vessel integrity, we indirectly evaluated interstitial tumor pressure in control and treated tumors. Histological assessment conducted seven days post-histotripsy demonstrated a significant reduction in tumor cell number per unit area in regions outside the treated zone, indicative of decreased tumor cell density (**Fig. S3A–D**). Seven days after histotripsy, blood vessel diameter outside the treated zone in B16F10 tumors increased approximately two-fold (from 0.007 mm to 0.014 mm) in targeted tumors, while off-target tumors exhibited a 1.6-fold increase in blood vessel diameter (averaged 0.011 mm) (**Fig. S3E–H**; **Table 5**). Similar patterns were noted across different tumor types and in both mouse and rat models (**Fig. S3I–K**; **Table 5**). Collectively, these findings suggest that partial histotripsy leads to a significant reduction in tumor cell density and a concomitant increase in blood vessel diameter within the TME, consistent with decreased interstitial tumor pressure. To further analyze blood vessel integrity, tomato lectin, which binds to sugar residues on endothelial cells, was introduced by tail vein injection in the bilateral flank mT4 model. Histotripsy-targeted tumors displayed significantly higher lectin perfusion than controls; contralateral off-target tumors exhibited non-significant increases in lectin perfusion compared to controls (**Fig. S4A–B**).

### Histotripsy-induced vascular normalization is antigen-specific

To investigate whether tumor vascular changes and normalization are immune-dependent, a hybrid tumor model was established by inoculating mice with Hepa1-6 tumors on one flank and B16F10 melanoma tumors on the opposite flank and treating only the Hepa1-6 tumor with histotripsy (**Fig. 3A**). Contralateral off-target B16F10 tumors were subjected to CD31 immunofluorescence for vasculature morphometry and intratumoral vasculature density analysis after sham and histotripsy treatment (**Fig. 3B**). Our results showed no reduction in most morphometric parameters in the discordant tumor following histotripsy, with total length of isolated vascular branches paradoxically increasing after histotripsy (**Fig. 3C**). Notably, there was still a significant decrease in total vessel density observed in the discordant tumor after contralateral histotripsy, suggesting that some TME effects of histotripsy could be antigen-non-specific (**Fig. 3D**). Overall, these findings suggest that the ability of histotripsy to induce off-target vascular normalization may be largely immune-mediated.

**Figure 3.**
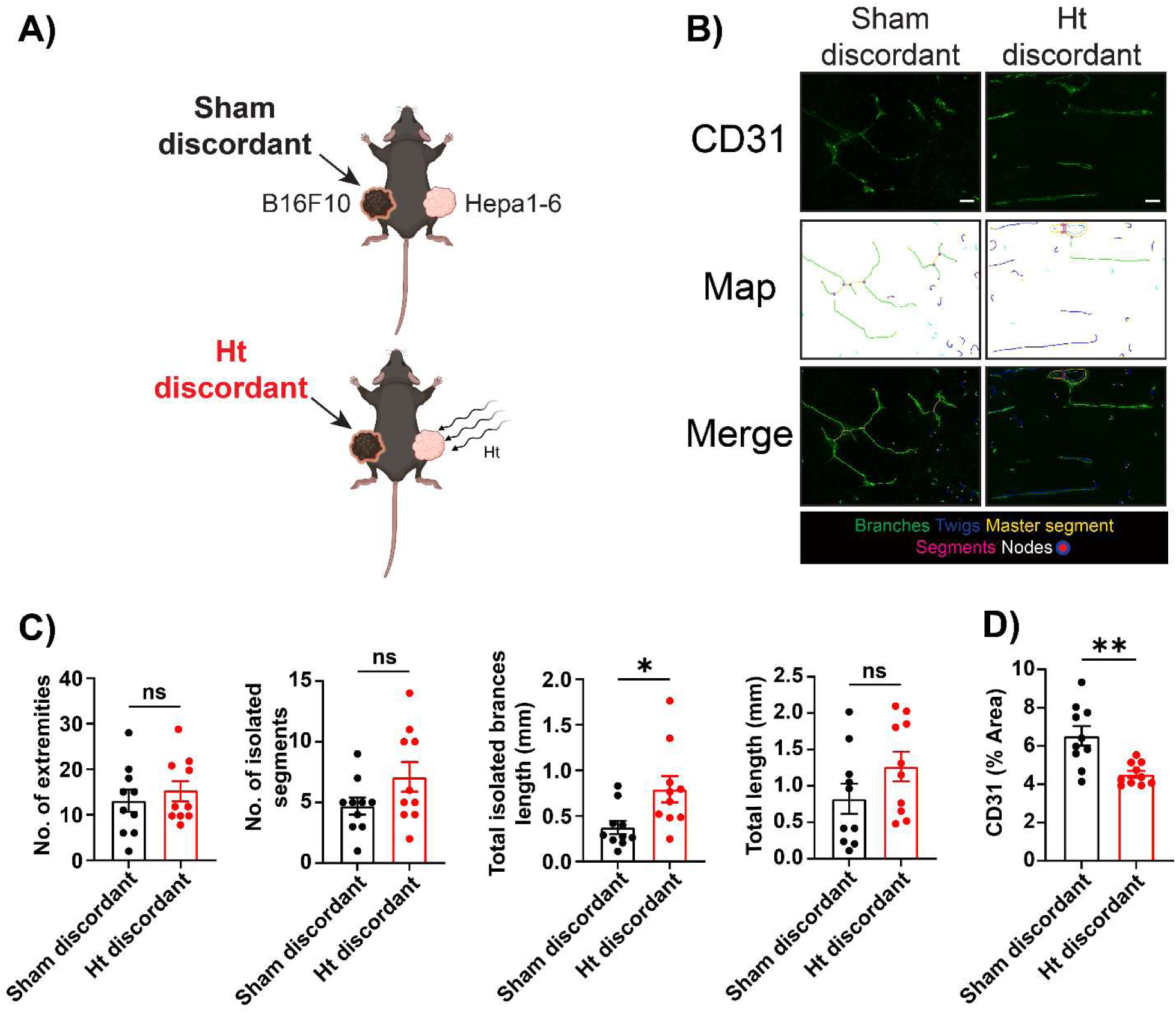
Histotripsy-induced vascular normalization is antigen-specific. (**A**) Hybrid model where Hepa1-6 tumors were inoculated on the right, and B16F10 tumors were inoculated on the left flank of mice. Histotripsy treatment was delivered on the right flank. (**B**) Tumor endothelium (CD31) staining of control (Sham discordant) and non-targeted contralateral (Ht discordant) tumors. Scale bars represent 50µm. (**C**) Quantification of different angiogenic features of the hybrid tumors. (**D**) Quantification of total vessel density in sham and histotripsy-treated discordant tumors (ns = not significant; *p < 0.05; **p < 0.01; error bars represent standard error of the mean (SEM)).

### CD8+ T cells are necessary for histotripsy-induced vascular normalization

Given our and other groups’ findings that histotripsy results in increased immune cell infiltration (*30–32*) and the mounting evidence for the crosstalk between CD8+ T cells and vascular remodeling (*33–35*), we assessed the relationship between CD8+ T cells and the presence of different features of angiogenesis in histotripsy-treated tumors. Our analysis identified a negative correlation between the presence of CD8+ T cells and angiogenic features, supporting the notion that CD8+ T cells regulate tumor vasculature, possibly by inducing endothelial cell apoptosis of immature tumor vessels by releasing interferon-γ (IFN-γ) as previously noted (*34–35*) (**Fig. 4A**). We validated the dependence of histotripsy-induced vascular normalization on CD8+ T cells by treating CD8-knockout C57BL/6 mice bearing bilateral B16F10 tumors with unilateral sham or partial histotripsy. We observed no significant differences in angiogenic features and total vessel density between control and histotripsy-treated tumors in mice lacking CD8+ T cells (**Fig. 4B–D**). Similarly, pericyte coverage, vascular stability, and leakiness markers revealed no differences following histotripsy treatment in the CD8-knockout mice (**Fig. 4E**; **Fig. S5A–E**). To further understand the role of CD8+ T cells in vascular normalization, we stimulated CD8+ T cells by administering anti-CTLA-4 mAb with or without histotripsy. Anti-CTLA-4 monotherapy significantly reduced angiogenic features and total vessel density, although to a lesser extent than histotripsy; combinatorial treatment showed an almost additive effect (**Fig. 4F–H**). Taken together, CD8+ T cells appear to be necessary to induce vascular normalization in histotripsy-treated tumors.

**Figure 4.**
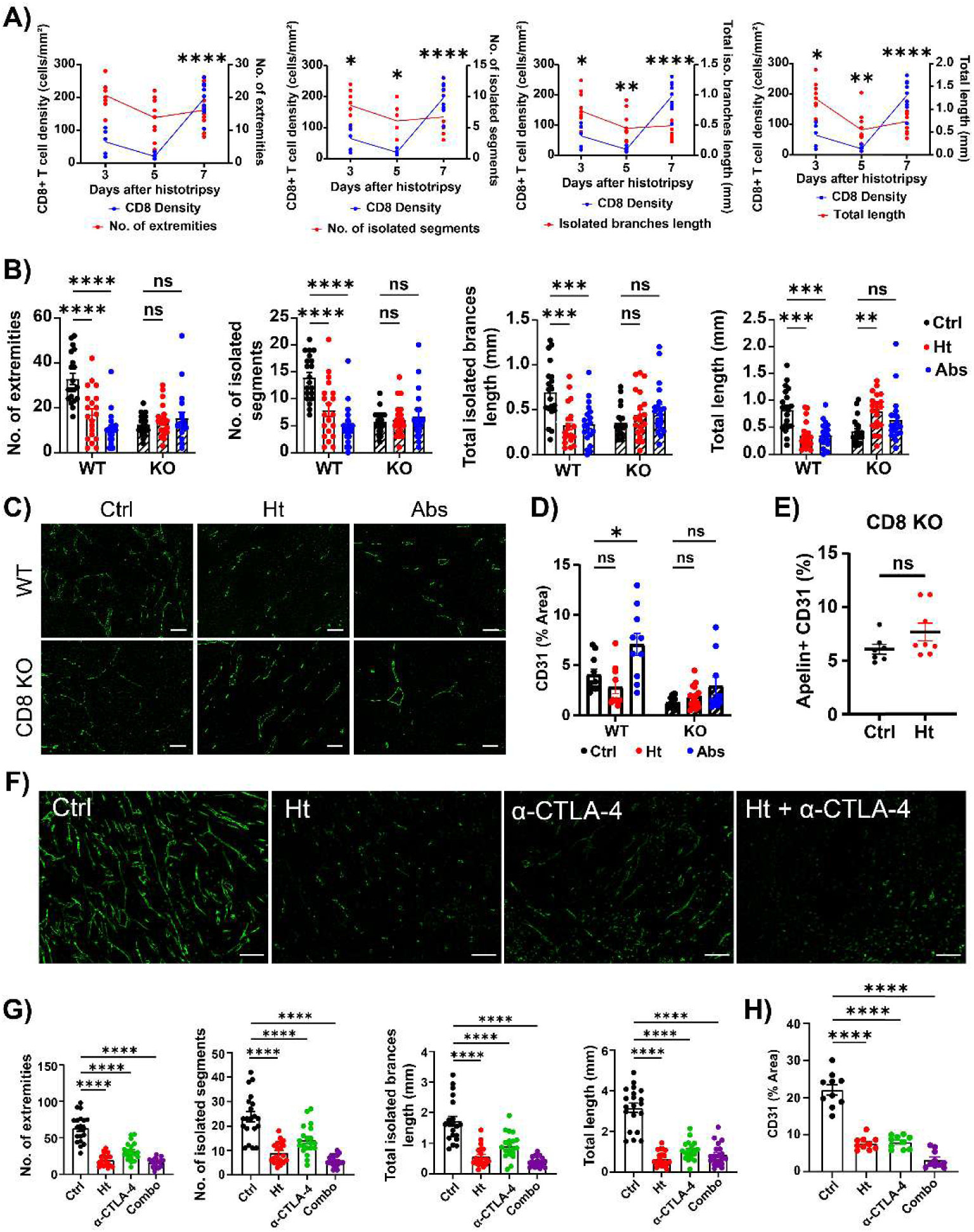
Vascular normalization is partially dependent on CD8+ T cells. (**A**) Graphical representation of different angiogenic features and CD8+ T cell infiltration over time in histotripsy-treated melanoma tumors. (**B**) Quantification of angiogenic features in WT and CD8-knockout (KO) mice. (**C**) IF staining of tumor endothelium (CD31) of wild-type (WT) and CD8 KO mice that were treated with sham or partial histotripsy. Scale bars represent 50µm. (**D**) Quantification of total vessel density in WT and CD8 KO mice subjected to sham (Ctrl) and partial histotripsy (Ht and Abs). (**E**) Quantification of Apelin+ endothelium in control (Ctrl) and treated (Ht) mice lacking CD8+ T cells. Representative images shown in **Fig. S5A**. (**F**) IF staining of tumor endothelium with CD31 in control (Ctrl), histotripsy-treated (Ht), anti-CTLA-4-treated (α-CTLA-4), and combinatorial (Ht + α-CTLA-4) treatment on Hepa1-6 tumor bearing mice. Scale bars represent 50µm. (**G**) Quantification of different features of angiogenesis from (**F**). (**H**) Quantification of total vessel density from (**F**) (ns = not significant; *p < 0.05; **p < 0.01; ***p < 0.001; ****p < 0.0001; error bars represent standard error of the mean (SEM)).

### CXCR3+CD8+ T cells are major contributors of vascular normalization

We have previously established that intratumoral CD8+ T cell infiltration after histotripsy is dependent on the CXCR3:CXCL10 signaling axis (*30*). Here, we used a B16F10 bilateral flank tumor model and administered neutralizing antibodies targeting CXCR3, effectively preventing CXCR3+CD8+ T cell infiltration. Immunofluorescence analysis showed that, without CXCR3+CD8+ T-cell infiltration, histotripsy does not alter vascular morphometry (**Fig. 5A–B**). Although there was a significant reduction in total vessel density in CXCR3-blocked mice, their baseline level of vessel density was almost doubled compared to the group that received IgG (**Fig. 5C**). As expected, we did not observe significant pericyte recruitment in histotripsy-treated mice that received blocking CXCR3 antibodies (**Fig. 5D–E**). In line with our observation, there was no difference in vasculature stability and leakiness as revealed by desmin and fibrin quantification, respectively, in CXCR3-blocked mice (**Fig. 5F–H**). Contralateral off-target tumors of mice that received anti-CXCR3 mAb showed significant differences in several features of angiogenesis and vascular stability, suggesting that the vasculature of the contralateral tumors could be mediated by cell types that do not rely on the CXCR3 signaling axis (**Fig. 5B**, **G**). These findings confirm that vascular normalization in histotripsy-treated tumors could be indirectly mediated by the CXCR3 signaling axis, likely via its role in facilitating CD8+ T cell recruitment to the tumor parenchyma.

**Figure 5.**
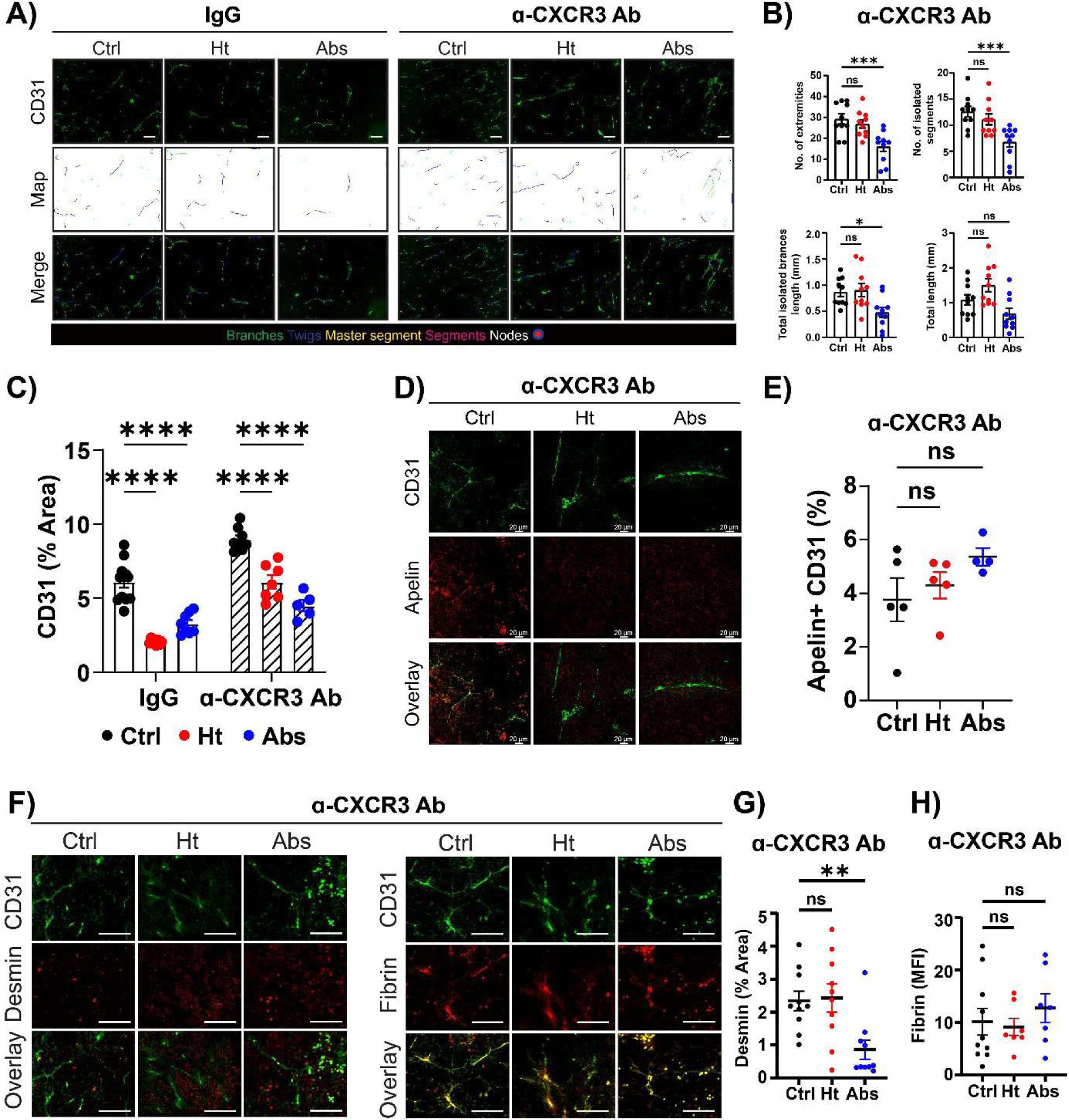
Histotripsy-induced vascular normalization is immune-cell mediated. (**A**) Tumor endothelium staining of IgG-treated and CXCR3-neutralized mice in control and histotripsy-treated groups. Scale bars represent 50µm. (**B**) Quantification of different features of angiogenesis from (**A**). (**C**) Quantification of total vessel density in control and histotripsy-treated mice subjected to IgG administration or neutralizing antibodies targeting CXCR3. (**D**) Co-IF staining with tumor endothelium (CD31) and pericyte (Apelin) in mice that received CXCR3 neutralizing antibodies. (**E**) Quantification of Apelin+ endothelium from (**D**). (**F**) IF staining of endothelial desmin and fibrin in mice treated with CXCR3 neutralizing antibodies. Scale bars represent 50µm. (**G–H**) Quantification of desmin and fibrin, respectively, from (**F**) (ns = not significant; *p < 0.05; **p < 0.01; ***p < 0.001; ****p < 0.0001; error bars represent standard error of the mean (SEM)).

### Histotripsy impacts the tumor vasculature by modulating the CXCR4 and Angiopoietin pathways

Previous studies have revealed that histotripsy abrogates hypoxia in murine models of melanoma and neuroblastoma (*30,36*). To provide mechanistic insights into what drives vascular normalization following histotripsy, we assessed several hypoxia-responsive proteins known to play roles in angiogenesis. First, we quantified endothelial CXCR4 expression in our different tumor types and found that its degree of post-histotripsy reduction is tumor-type specific (**Fig. 6A–D**). B16F10 tumors had a four-fold decrease in endothelial CXCR4 expression after histotripsy in both treated and contralateral off-target tumors; while Hepa1-6 tumors did not show significant reduction, we still saw a downward trend in endothelial CXCR4 expression after histotripsy (**Table 6**). Congruent with the melanoma model, histotripsy-treated pancreatic mT4 tumors showed a four-fold reduction in endothelial CXCR4 expression, with contralateral off-target tumors showing a decrease comparable to targeted tumors (**Table 6**). Secondly, we utilized multi-color immunofluorescence to stain for Angpt1 and Angpt2 expression on tumor endothelial cells (**Fig. 6E**). In B16F10 and Hepa1-6 tumors, histotripsy treatment increased Angpt1:Angpt2 ratio by two-fold, compatible with vessel maturity and stabilization (**Table 7**). Strikingly, mT4 tumors did not exhibit statistically significant differences in Angpt1:Angpt2 ratio after histotripsy, although we still observed a 1.8-fold increase compared to the control (**Fig. 6F–H**; **Table 7**). Overall, while we have not elucidated the specific factors that directly activate these pathways, our data suggest that histotripsy may modulate the CXCR4 and angiopoietin pathways to induce vascular normalization.

**Figure 6.**
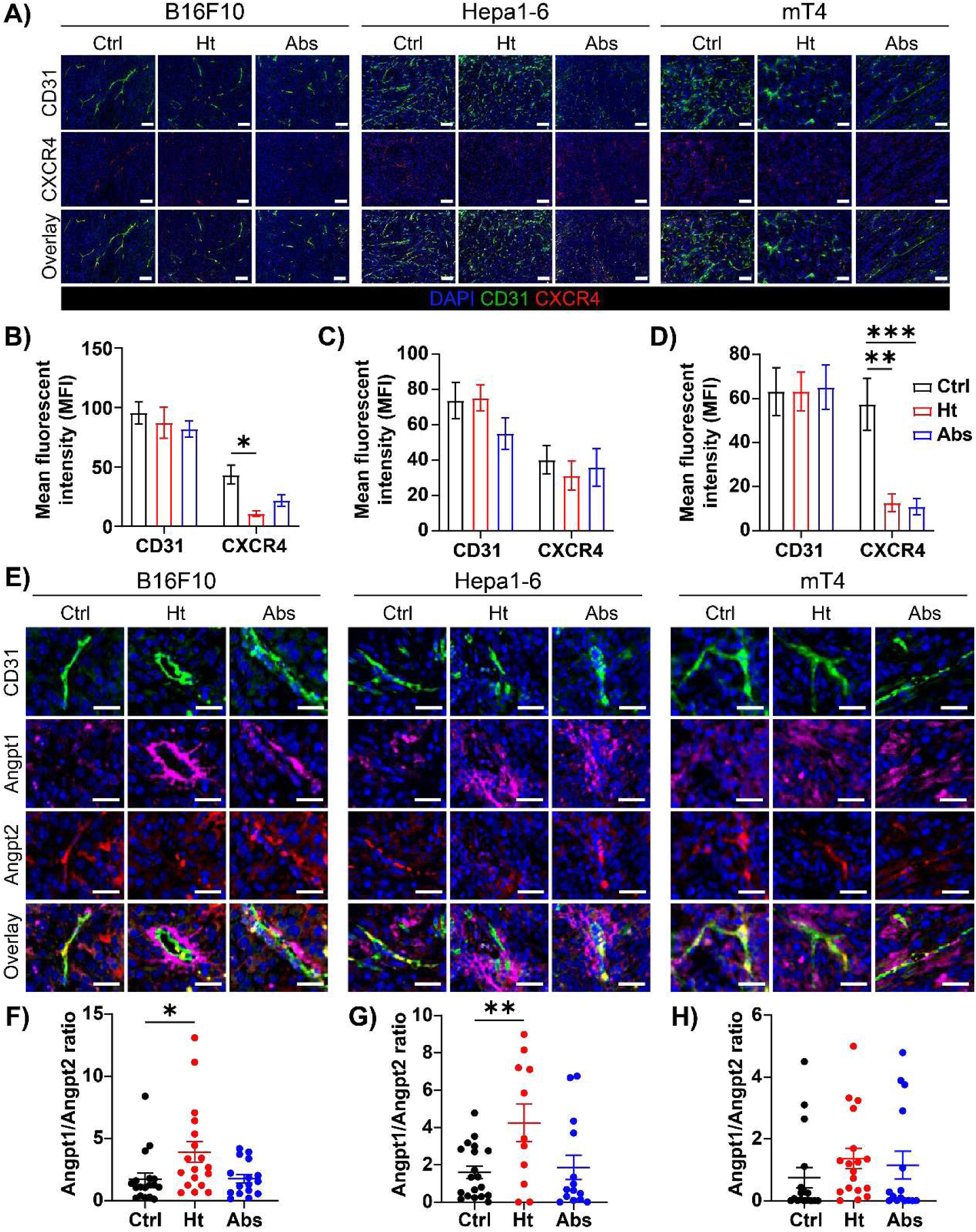
Histotripsy impacts the tumor vasculature by modulating the CXCR4 and Angiopoietin pathways. (**A**) IF staining of endothelial CXCR4 in B16F10, Hepa1-6, and mT4 tumors. Scale bars represent 50µm. (**B–C**) MFI of endothelial CXCR4 expression from (**A**). (**E**) Multicolor IF staining of CD31, angiopoietin-1, and angiopoietin-2 in control and histotripsy-treated tumors. Scale bars represent 20µm. (**F–H**) Quantification of angiopoietin-1 and angiopoietin-2 ratio from (**E**) (ns = not significant; *p < 0.05; **p < 0.01; ***p < 0.001; error bars represent standard error of the mean (SEM)).

### Histotripsy enhances intratumoral drug delivery

To investigate whether histotripsy-induced vascular remodeling is associated with improved immune cell infiltration, we analyzed intratumoral CD8+ T cell proximity with respect to tumor blood vessels. Strikingly, our analysis showed a significant increase in CD8+ T cell migration after histotripsy in B16F10 and mT4 tumors, and to a lesser extent in Hepa1-6 tumors (**Fig. 7A–C**). We then evaluated the effect of histotripsy on chemotherapeutic and immunotherapeutic drug delivery by injecting doxorubicin or anti-CTLA-4 mAb into the tail vein and measuring doxorubicin autofluorescence (**Fig. 7D–G**) or mAb presence (**Fig. 7H–I**) in control tumors and in treated and off-target tumors after histotripsy. We found that histotripsy-treated tumors were more accessible to both chemotherapeutic drugs and monoclonal antibodies compared to control tumors. While doxorubicin delivery into B16F10 and mT4 off-target total tumors did not show a significant increase, we did observe significant drug penetration into the periphery of the off-target contralateral tumors after histotripsy (**Fig. 7D–G**). Increased immune cell infiltration and enhanced drug delivery following treatment indicate that histotripsy has the capability to promote functional tumor blood vessels, potentially alleviating the immunosuppressive TME and increasing the efficacy of conventional therapies.

**Figure 7.**
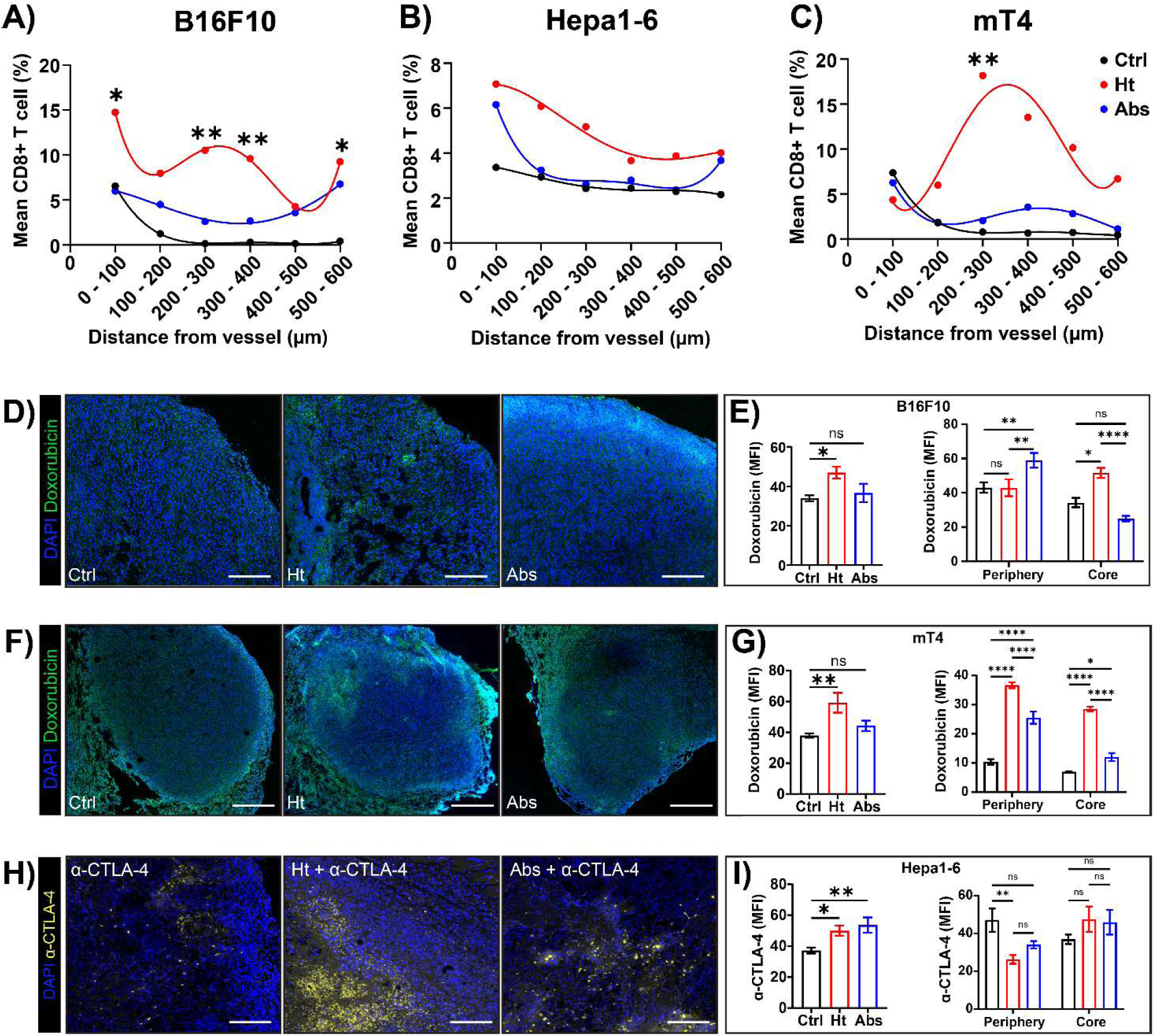
Histotripsy impacts intratumoral drug delivery. (**A-C**) Quantification of CD8+ T cell density relative to the distance from the vessel in control and histotripsy-treated B16F10, Hepa1-6, and mT4 tumors respectively. (**D**) Doxorubicin delivery to B16F10 tumors via tail vein injection 7 days after histotripsy. Scale bars represent 1mm. (**E**) Quantification of doxorubicin delivery into the whole tumor, including comparison between tumor periphery and core, from (**D**) using MFI. (**F**) Doxorubicin delivery to mT4 tumors via tail vein injection 10 days after histotripsy. Scale bars represent 1mm. (**G**) Quantitation of doxorubicin delivery from (**F**) using MFI. (**H**) Staining of anti-CTLA-4 in Hepa1-6 tumors treated with either monotherapy (α-CTLA-4) or combination therapy (Ht + α-CTLA-4). Scale bars represent 100µm. (**I**) Quantitation of anti-CTLA-4 delivery to Hepa1-6 tumors using MFI (ns = not significant; *p < 0.05; **p < 0.01; ***p < 0.001; ****p < 0.0001; error bars represent standard error of the mean (SEM)).

## DISCUSSION

Solid tumors depend on continuous blood supply for growth and metastasis that is achieved through VEGF-driven vascular proliferation and angiogenesis (*3,5,10*). By targeting this process, clinical antiangiogenic therapy aims to prune abnormal vessels to control tumor growth and potentially improve drug delivery via vascular normalization (*21,22,37*). In this study, we found that histotripsy results in significant reductions in total vessel density across different tumor types. Of note, melanoma and HCC are known for their hypervascular nature that promotes aggressive tumor growth and metastasis, while pancreatic adenocarcinoma (PDAC) is inherently hypovascular, resulting in poor perfusion and inefficient drug uptake (*38–40*). Unlike the significant remodeling of vascular anatomy seen in B16F10 (melanoma) and Hepa1-6 (HCC) models, the remaining blood vessels in mT4 (PDAC) tumors retained their angiogenic features despite a significant reduction in total vessel density. In spite of these differences, we observed evidence of enhanced tumor perfusion as measured by tomato lectin or doxorubicin or therapeutic antibody delivery in all three tumor types. Taken together, our data indicate that histotripsy induces favorable vascular remodeling that is largely independent of intrinsic tumor vascularity status. High tumor cell density causes increased intratumoral pressure through the solid stress generated by rapidly proliferating cells, resulting in tumor blood vessel constriction. Increased pressure also results from interstitial fluid buildup as basement membrane discontinuity and poor pericyte coverage of blood vessel endothelium leads to leaky tumor vasculature (*16,17,41*). Here, we observed a significant decrease in tumor cell density accompanied by increased blood vessel diameter after histotripsy, suggesting a decrease in tumor pressure following histotripsy—a phenomenon that could be verified by direct measurement of tumor interstitial fluid pressure in future studies.

Vascular normalization is a highly orchestrated process that relies on restoring the balance between pro- and anti-angiogenic factors within the TME (*21,37,42*). Our previous study revealed that VEGF expression is abrogated following histotripsy (*30*); in the current study, we therefore measured the angiogenic potential of tumor blood vessels by quantifying morphological features of angiogenesis including number of extremities, number of isolated branches, total isolated branches length, and total branches length, all of which were significantly reduced following histotripsy in our melanoma and hepatocellular carcinoma models. In addition to characterizing morphological changes, we also interrogated changes in vascular physiology by assessing the integrity of the tumor endothelium using markers for pericytes such as apelin and desmin, in which their presence reflects vessel stability and maturation. We show that in histotripsy-treated mice, tumor blood vessels exhibited increased expression of apelin and desmin, indicative of vessel stabilization and maturation. We also identified increased perfusion measured by lectin probe and reduced vascular leakiness determined by fibrin extravasation following histotripsy. Importantly, these changes were seen not only in directly treated tumors but also in off-target tumors, highlighting an abscopal or systemic effect on tumor vasculature.

Immune-vascular crosstalk is fundamental to this process, as immune cells contribute to the remodeling of hyperangiogenic tumor blood vessels (*33–35*), and normalized vasculature supports immune cell infiltration and amplifies anti-tumor immunity (*34,37,43*). Strategies combining angiogenesis inhibitors with immune checkpoint inhibitors (ICIs) are being used in an effort to normalize tumor vasculature and facilitate cytotoxic lymphocyte entry into the TME, thereby potentiating the efficacy of immunotherapy (*37,42,43,44*). The rationale for these combinatorial strategies is strengthened by recent studies highlighting the immune-driven nature of vascular normalization, as cytotoxic immune cells such as CD8+ T cells release angiostatic factors to suppress abnormal angiogenesis, promote vessel maturation, and upregulate endothelial adhesion molecules (*33–35*). Previous studies have demonstrated that CD8+ T cells influence tumor vasculature normalization and pruning by inducing endothelial cell apoptosis of immature tumor vessels through the release of IFN-γ (*33–35*). Given the link between vascular normalization and antigen specificity, we hypothesized that CD8+ T cells are a major contributor to the vascular effects of histotripsy. Indeed, vascular normalization was impaired in histotripsy-treated mice with genetic knockout of CD8+ T cells, while re-invigoration of CD8+ T cells with ICI enhanced this effect. Our previous data revealed that the CXCL9/10/11:CXCR3 axis is imperative in promoting CD8+ T cell infiltration into the tumor following histotripsy (*30*). As expected, blocking the CXCL9/10/11:CXCR3 pathway diminished the effects of histotripsy on vessel remodeling, further supporting the role of CD8+ T cells in histotripsy-induced vascular normalization. Importantly, the antigenic specificity of this abscopal vascular remodeling and normalization after histotripsy seen in our hybrid model of antigenically discordant tumors reinforces the concept of immune-mediated vascular normalization. While angiogenic features in mice that received anti-CXCR3 neutralizing antibodies post-histotripsy remained unchanged, total vessel density was significantly decreased in both treated and contralateral off-target tumors, suggesting that vascular remodeling may not be entirely dependent on this axis. The lack of vascular remodeling of treated tumors following CXCR3 blockade suggests that CXCR3+CD8+ T cells may be necessary to drive the reorganization of vessel structure following histotripsy. This is compatible with the clinical strategy of using antiangiogenic agents to induce vascular normalization and ICIs to activate cytotoxic T cells. This strategy has proven to be successful in prolonging progression-free survival (PFS) in HCC and renal cell carcinoma (*25,44,45*). Of note, despite their clinical benefits, antiangiogenic agents are not universally effective, and the dosage and timing of administration can significantly affect therapeutic outcomes (*23,24,26*).

One of the ways solid tumors adapt to the hypoxic TME is by upregulating HREs including HIF-1, VEGF, and the CXCL12 receptor, CXCR4 (*5,7,9*). We previously demonstrated that histotripsy abrogates HIF-1 and VEGF in murine models of melanoma (*30*); in the current study, we explored whether post-histotripsy vascular remodeling is mediated by CXCR4, given its role in promoting blood vessel sprouting (*10*). Our results show that endothelial CXCR4 is significantly diminished in our melanoma and PDAC mouse models but only modestly in HCC, revealing distinct CXCR4 modulation in different tumor types. To further distinguish mature vessels from dysfunctional ones, we measured the ratio of Angpt1 and Angpt2 on the tumor endothelium of sham and histotripsy-treated mice. Our observation that vascular normalization is accompanied by an increased Angpt1:Angpt2 ratio following histotripsy is in line with previous observations wherein Angpt1 promotes vessel stabilization by restoring a hierarchical vasculature, while Angpt2 antagonizes this process (*11,13*). Thus, in our future studies, we will be modulating this pathway and assessing its impact on vascular normalization following histotripsy.

Based on our findings that histotripsy enhances blood vessel integrity, we tested CD8+ T-cell trafficking and drug delivery into the tumors. To this end, we quantified the percentage of CD8+ T cells, which is defined as the number of CD8+ T cells in selected areas of interest, relative to their distance from a tumor vessel. In control tumors, average CD8+ T-cell percentage decreased the farther we measured from the vessel, whereas in histotripsy-treated tumors, there were significant increases in average CD8+ T-cell density relative to their distance from a tumor vessel, suggesting enhanced CD8+ T cell migratory capacity. Alternatively, CD8+ T-cell infiltration could be influenced by hypoxia gradients in the tumor given our previous observation that CD8+ T cell infiltration is geographically higher in tumor regions where HIF-1α is downregulated (*30*). Moreover, we took advantage of the autofluorescent properties of the chemotherapeutic drug doxorubicin to measure intratumoral drug delivery following histotripsy. Our results show that tumors from histotripsy-treated mice had significant Doxorubicin uptake compared to untreated mice. We identified that increased drug delivery is not limited to chemotherapy, but also applies to monoclonal antibodies as demonstrated by enhanced anti-CTLA-4 penetration into the tumor following histotripsy. This observation addresses one of the limitations associated with antiangiogenic therapy whereby treatment with bevacizumab reportedly decreases antibody uptake, necessitating careful therapeutic strategies when using combinatorial treatment with the drug (*46*). Our findings suggest that enhanced drug delivery following histotripsy-induced vascular remodeling is not limited to drugs but also extends to larger proteins, demonstrating yet another clinical advantage over conventional antiangiogenic agents. Notably, we observed that significant anatomic and physiological alterations following histotripsy are necessary to enhance drug delivery into contralateral, off-target tumors, and that the addition of ICIs following histotripsy could further improve drug penetration into distant tumors as demonstrated by our Hepa1-6 model. Taken together, our data reveal that localized histotripsy treatment triggers an abscopal effect that could be potentiated by the addition of ICIs to improve drug delivery and facilitate control of distant tumors, which may have significant clinical implications in the treatment of metastatic tumors.

Due to histotripsy’s favorable safety profile and ability to stimulate systemic immune responses as observed in pre-clinical and early clinical trials prior to its FDA approval for the treatment of liver cancers (*30,31,32,47*), there are currently several ongoing clinical trials testing its feasibility in different types of solid cancers including primary renal tumors (NCT05820087) and pancreatic adenocarcinoma (NCT06282809). Here we demonstrate an additional advantage of histotripsy compared to other methods of ablation therapy, wherein localized treatment results in systemic vascular normalization as demonstrated by significant vascular remodeling in both treated and non-treated contralateral tumors. In this study, we provide proof of principle that histotripsy can normalize tumor vasculature, ultimately leading to enhanced drug delivery and tumor control.

## MATERIALS AND METHODS

### Study design

The objective of this study was to elucidate the mechanism by which histotripsy results in increased immune cell infiltration and enhanced drug delivery. We hypothesized that liquefaction of solid cancers following histotripsy leads to the reorganization of the tumor vasculature, improving drug delivery and potentiating antitumor immune responses. We treated several *in vivo* models of solid tumors with varying vascularity status, namely melanoma, hepatocellular carcinoma, and pancreatic adenocarcinoma to elucidate the extent of vascular remodeling and normalization that occurs following histotripsy. We performed co-immunofluorescence (co-IF) and microscopy studies to evaluate and quantitate markers for angiogenesis and vascular normalization. We assessed the role of the CXCR3 axis and CD8+ T cells by performing microscopic analyses in mice that received anti-CXCR3 neutralizing antibodies and CD8-knockout mice, respectively. Intratumoral uptake of doxorubicin and anti-CTLA-4 monoclonal antibodies were quantified to measure drug delivery in control and histotripsy-treated tumors.

### Cell lines

B16F10, Hepa1-6, and mT4 cell lines were purchased from ATCC and have been authenticated by the vendor. B16F10 was maintained in culture with RPMI-1640 medium (Gibco, Life Technologies, Grand Island, New York) with 10% fetal bovine serum (R&D Systems, Minneapolis, Minnesota) and 1% penicillin/streptomycin (Gibco, Life Technologies). Hepa1-6 and mT4 cells were maintained in DMEM (Gibco, Life Technologies) with 10% fetal bovine serum (HyClone, GE Healthcare Life Sciences, Marlborough, Massachusetts) and 1% penicillin/streptomycin (Gibco, Life Technologies). Rat-derived McA-RH7777 cells (ATCC® CRL-1601™) purchased from ATCC were cultivated in DMEM (Gibco, Life Technologies), 10% fetal bovine serum (Corning), and 1% penicillin/streptomycin (Gibco, Life Technologies). The cell lines were routinely checked for mycoplasma contamination, and only the first 10 passages were used for experiments.

### Animal studies

Male and female C57BL/6 mice aged 6–8 weeks old were purchased from Taconic (Hudson, New York) and housed and maintained in specific pathogen-free conditions. CD8 knockout mice were purchased from Jackson Laboratory (Bar Harbor, Maine) and maintained in specific pathogen-free conditions. Each experiment involved 4-10 mice per experimental group. Flank tumors were established by subcutaneous injection with 1x10^6^ cells of the appropriate cell line suspended in phosphate-buffered saline (PBS). mT4 tumors were suspended in 1:1 PBS:Matrigel prior to tumor inoculation. Flank tumors were monitored and measured with electronic calipers daily after tumor establishment, and tumor volumes were calculated according to the following formula: volume= 4/3 x π x (long dimension/2) x (short dimension/2) x (long dimension/3). Endpoint criteria for euthanasia included maximal tumor diameter >20 mm with concomitant >50% tumor ulceration. Mice were routinely monitored by body condition (BC) scoring. Mice with less than a score of BC2 were dropped from the study and humanely euthanized. The murine experiments were prospectively reviewed and approved by the VA Ann Arbor Healthcare and University of Michigan Animal Care and Use Committees. Male Sprague Dawley rats weighing 100-125 g were purchased from Taconic (Hudson, New York) and housed and maintained in specific pathogen-free conditions. Orthotopic liver tumors were established by injecting 10x10^6^ McA-RH7777 cells suspended in PBS and Matrigel in 1:1 ratio into the liver following laparotomy, as established previously (*48,49*). Tumors were monitored using ultrasound imaging, and endpoint criteria included maximal tumor diameter >20 mm. Study approval was obtained from the Institutional Animal Care and Use Committee at the University of Michigan.

### Histotripsy

Histotripsy treatment was performed using the 1MHz rodent therapy transducer. Approximately 60-80% tumor volume was targeted using a pulse repetition frequency (PRF) of 100Hz, and 50 – 100 pulses were delivered at each focal location to achieve an estimated −30 MPa peak negative pressure, monitored under real-time ultrasound guidance (*31*). The spacing between the targeted locations varied between individual mice from 0.5 – 0.6 mm in the x, y, and z dimensions. The spacing was adjusted to keep treatment times relatively the same. All procedures were performed following induction of isoflurane anesthesia. Sham treatment received isoflurane anesthesia alone. Prior to histotripsy, mice received Lidocaine (1mg/kg) for analgesia. Rat liver tumors were also targeted with the 1 MHz rodent therapy system, using a PRF of 100 Hz, with 100 pulses delivered at each location, with a spacing of 0.6 mm – 0.7 mm between individual locations. Carprofen (5 mg/kg) was administered subcutaneously prior to treatment for analgesia and inhaled isoflurane was used for anesthesia. Animals were randomized to control versus histotripsy treatment. Control treatments were delivered by placing animals in the same setup as the histotripsy group and administering sham exposures, without activation of the histotripsy device. Tumors of sham- and histotripsy-treated animals were harvested after euthanasia for morphometric assessment of tumor vasculature.

### Antibody and drug treatment

Selected mice received intraperitoneal (i.p.) injections of 200 µg anti-CXCR3 antibody (clone CXCR3-173, BioXcell, Lebanon, New Hampshire) or control IgG (clone BE0091, BioXcell) twice a week until the conclusion of the experiments. Doxorubicin solution was prepared as per manufacturer’s protocol (MilliporeSigma, Burlington, Massachusetts). In brief, 100 µL of doxorubicin was administered via tail vein injection into each mouse one hour before tumor harvest. Lycopersicon Esculentum (tomato) Lectin DyLight 649 (Vector Laboratories, Newark, California) was injected via tail vein injection into selected mice one hour before tumor harvest. PBS was injected into mice to act as a negative control in both doxorubicin and lectin experiments.

### Immunofluorescence and analysis

For formalin-fixed paraffin-embedded (FFPE) tissues, harvested tumors were washed with PBS and preserved in 10% buffered formalin for 24-72 hours and in 70% ethanol until paraffin embedding. For fresh frozen tissues, tumors were washed with PBS and embedded into casts with Optimal Cutting Temperature (OCT) compound (Sakura Finetek, Torrance, California) and sectioned with a Leica CM1520 cryostat. FFPE sections were deparaffinized in xylene, then rehydrated by sequential transfer to decreasing concentrations of ethanol followed by distilled water. Antigen retrieval was performed using citrate buffer (ThermoFisher Scientific, Waltham, Massachusetts) at 70-90°C for 15 minutes. Samples were washed with PBS and blocked in 10% bovine serum albumin (ThermoFisher Scientific) for 1 hr at RT. Frozen sections were fixed with 3.5% PFA (Electron Microscopy Sciences, Hatfield, Pennsylvania) and permeabilized with 0.5% TritonX-100 (ThermoFisher Scientific) before blocking. Samples were incubated with primary antibody cocktail, diluted at 1:100 concentration with universal antibody dilution solution (Sigma Aldrich, St. Louis, Missouri) at 4°C overnight. Primary antibodies used were goat anti-CD31 (R&D Systems), rabbit anti-apelin (ThermoFisher Scientific), rabbit anti-desmin (Sigma-Aldrich, Burlington, Massachusetts), mouse anti-fibrin (clone 59D8) (Sigma-Aldrich), rabbit anti-CXCR4 (clone EPUMBR3) (Abcam, Cambridge, England, UK), rabbit anti-angiopoietin 1 (Abcam), and mouse anti-angiopoietin 2 (R&D Systems). Samples were counterstained with a secondary antibody cocktail diluted at 1:200 concentration with universal antibody dilution solution (Sigma-Aldrich), and incubated at 37°C for 60 min. Secondary antibodies used were donkey anti-goat Alexa Fluor Plus 488, donkey anti-mouse Alexa Fluor 555, donkey anti-rabbit Alexa Fluor 647, and chicken anti-rabbit Alexa Fluor 647 (Invitrogen, Carlsbad, California). Samples were mounted with mounting media containing DAPI (Vector Laboratories). Rat FFPE samples were submitted to ULAM Pathology Core for sectioning and CD31 staining. Slides were visualized using Keyence BZ-X810 fluorescence microscope (Keyence, Osaka, Japan) under 20X objective with 1X digital zoom and Leica Stellaris 5 inverted confocal microscope (Leica Microsystems, Wetzlar, Germany) under 40X objective with 1X digital zoom. Spatial intensity of staining was performed in ImageJ (National Institutes of Health, Bethesda, Maryland). Angiogenesis Analyzer plug-in in ImageJ was utilized to quantify morphometric changes in the tumor vasculature. A minimum of 10 fields of view from at least two independent experiments were considered for intensity and statistical analysis.

## Statistical analysis

Statistical analysis was performed using GraphPad Prism software (GraphPad Software, San Diego, California). The difference between means of unpaired samples was performed using one-way analysis of variance (ANOVA) with Tukey’s multiple comparison test or a two-tailed t-test, unless otherwise stated. Tumor growth kinetics and grouped datasets were compared using two-way ANOVA with Tukey’s multiple comparison test, comparing columns within each row, unless otherwise stated. Survival curves were analyzed with the logrank test. Statistical significance was defined as p<0.05. Numbers of mice per experiment are noted in the figure legends.

## Acknowledgments

We thank members of the Xu Lab for their valuable assistance with this project. This work was performed with instrumentation at the VA Ann Arbor Healthcare System Research Services. The authors wish to acknowledge the support of the University of Michigan Pathology Core for rat histology and immunohistochemistry staining. The content is solely the responsibility of the authors and does not represent the views of the Department of Veterans Affairs or the United States Government or the National Institutes of Health. The authors also wish to acknowledge the University of Michigan Undergraduate Research Opportunity Program for their support of C.K., N.G., and K.B., and the University of Michigan Rackham Merit Fellowship to support H.Q.

## Author contributions

Conceptualization: AG, CSC

Methodology: HQ, BS, HK, MG, JL, AS, KB, CK, AB, TW, ZX, AG

Investigation: HQ, AG, CSC Visualization: HQ, BS, JL

Funding acquisition: ZX, CSC, AG Project administration: AG, CSC Supervision: AG, CSC

Writing – original draft: HQ, AG, CSC

Writing – review & editing: HQ, AG, TW, CSC

## Data and materials availability

All data associated with this study are present in the paper or the Supplementary Materials. All materials used or generated in this study are commercially available or will be supplied upon reasonable request.

