## Supplemental Figures and Tables for "Histotripsy-induced vascular remodeling in the tumor microenvironment enhances local and off-target intratumoral drug delivery"

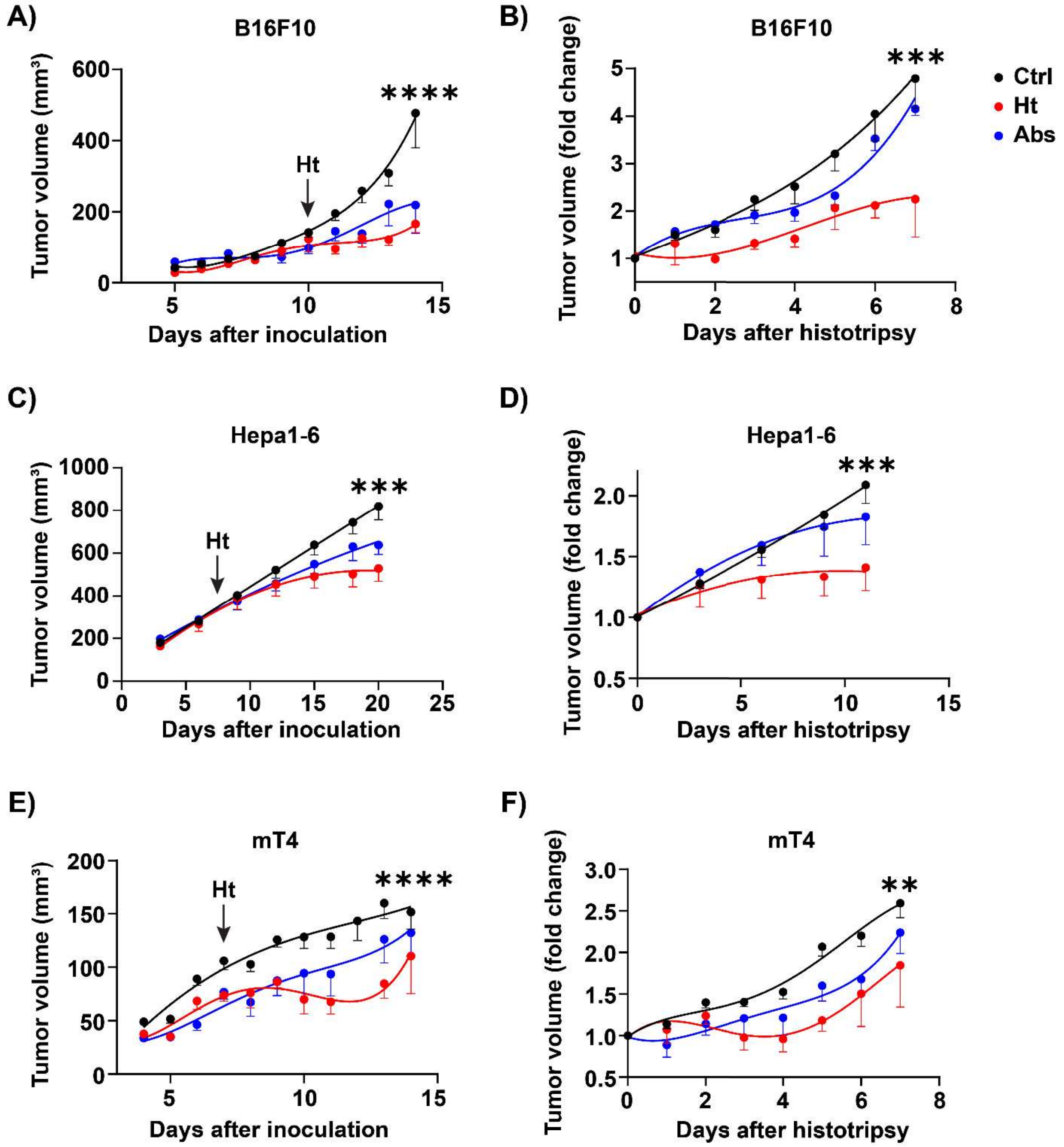


**Supplemental Figure S1. Histotripsy decreases tumor growth kinetics of different tumor-types.** (**A–B**) Tumor growth kinetics of mice bearing B16F10 tumors subjected to sham or partial histotripsy. (**C–D**) Tumor growth kinetics of mice bearing Hepa1-6 tumors treated with sham or partial histotripsy. (**E–F**) Tumor growth kinetics of mice bearing mT4 tumors subjected to sham or partial histotripsy (ns = not significant; $^{**}p < 0.01$; $^{***}p < 0.001$; $^{****}p < 0.0001$; error bars represent standard error of the mean (SEM)).

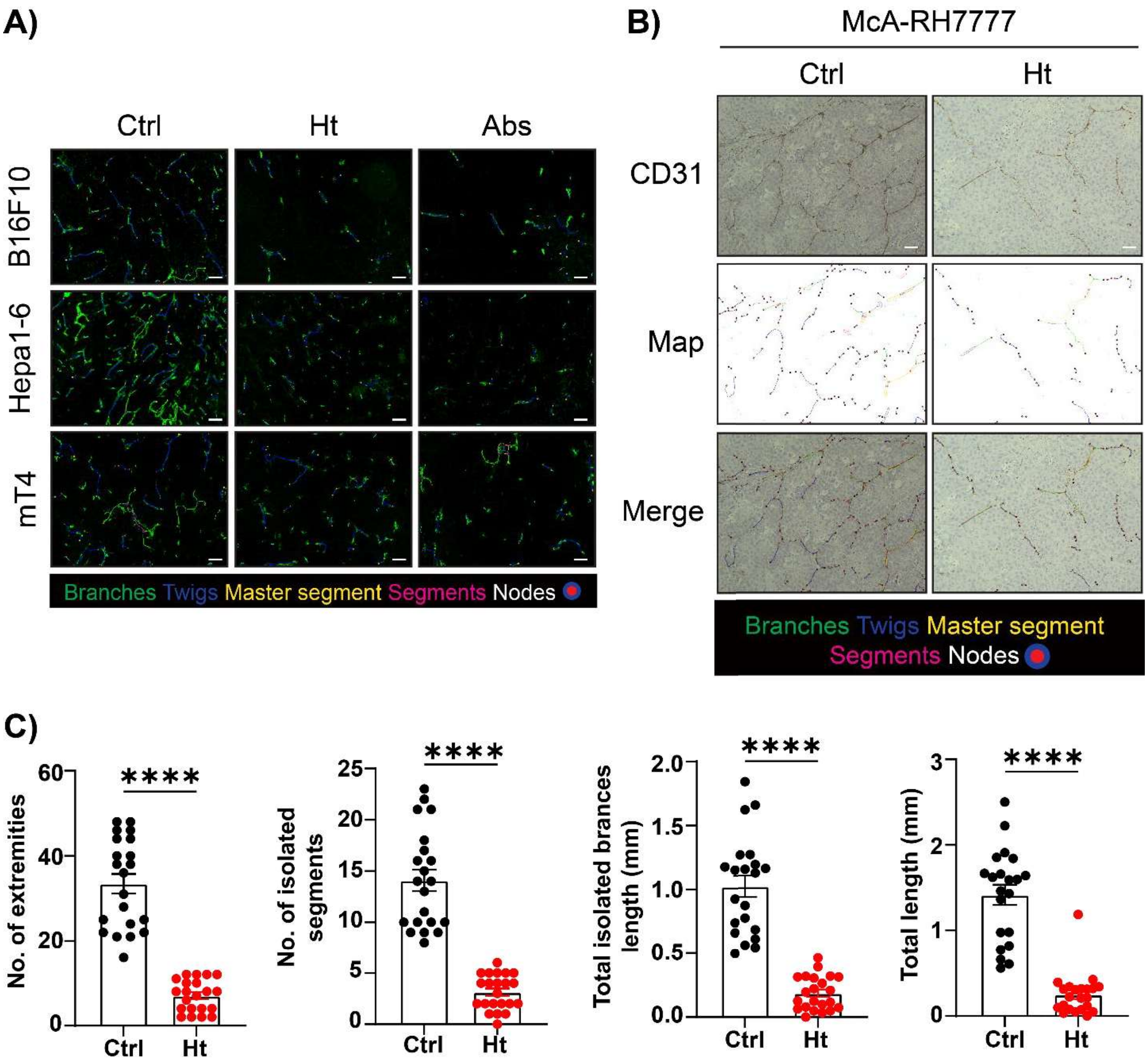


**Supplemental Figure S2. Histotripsy alters the vascular architecture of several tumor types.** (**A**) Overlay of tumor endothelium staining with different features of angiogenesis from **Fig. 1**. Scale bars represent 50μm. (**B**) Tumor endothelium (CD31) immunohistochemical staining of McA-RH7777 tumors, with maps showing different features of angiogenesis: branches, twigs, masters segment, segments, and nodes. Scale bars represent 100μm. (**C**) Quantification of four different angiogenic features from (**B**) (ns = not significant; ****p < 0.0001; error bars represent standard error of the mean (SEM)).

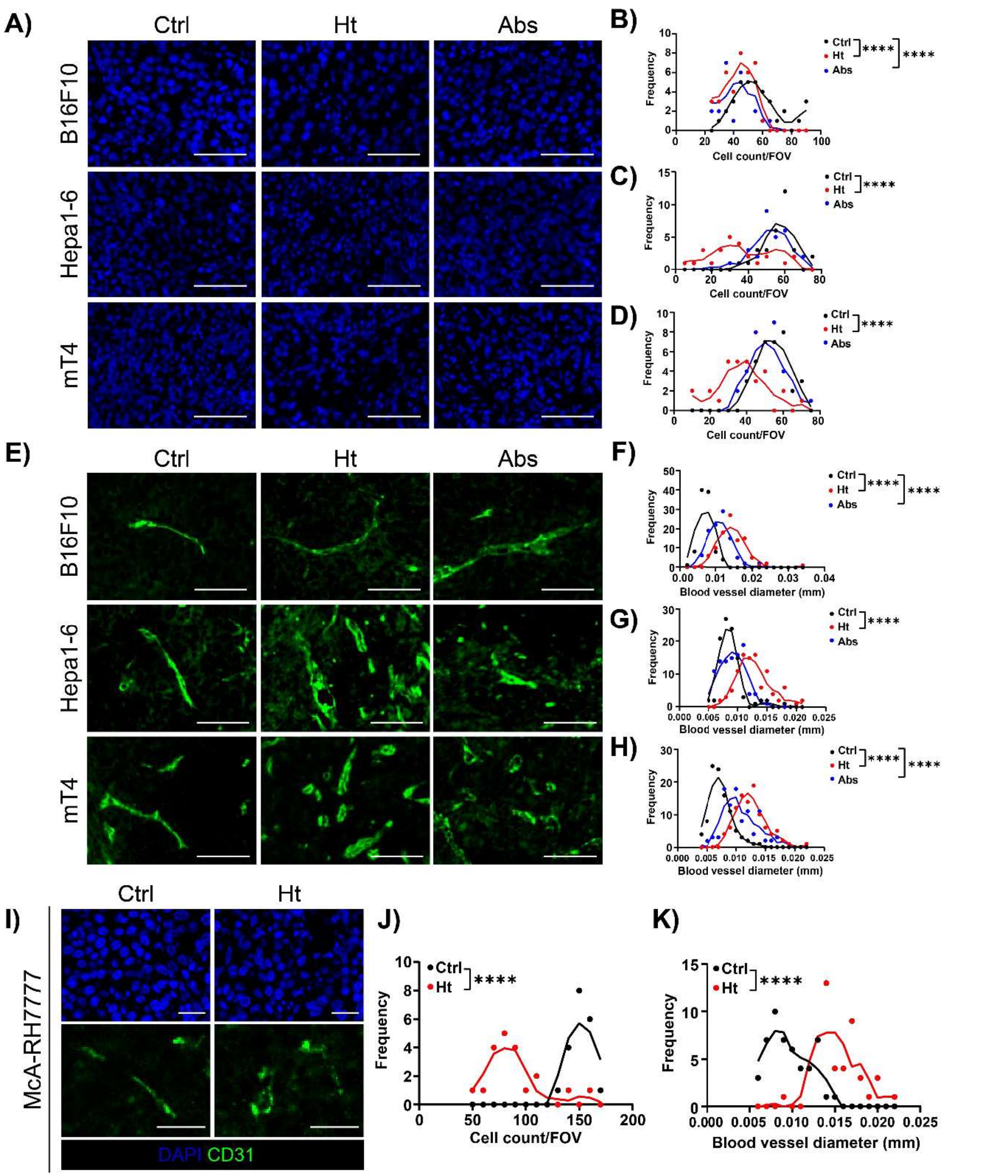


**Supplemental Figure S3. Decrease in intratumoral cell density and increase in blood vessel diameter after histotripsy.** (**A**) DAPI staining of B16F10, Hepa1-6, and mT4 tumors from untreated (Ctrl), treated (Ht), and non-treated contralateral (Abs) mice. Scale bars represent 50µm. (**B–D**) Frequency of cells counted from ≥100 field of view (FOV) in different tumor-types of sham and treated mice. (**E**) IF staining of B16F10, Hepa1-6, and mT4 tumor blood vessel using anti-CD31. Scale bars represent 50µm. (**F–H**) Quantification of blood vessel diameter from (**E**). (**I**) DAPI and CD31 staining of McA-RH7777 tumors from sham (Ctrl) and histotripsy-treated (Ht) rats. Scale bars represent 20µm. (**J**) Frequency of cells counted from ≥100 FOV from (**I**). (**K**) Quantification of blood vessel diameter from (**I**) (ns = not significant; ****$p < 0.0001$; error bars represent standard error of the mean (SEM)).

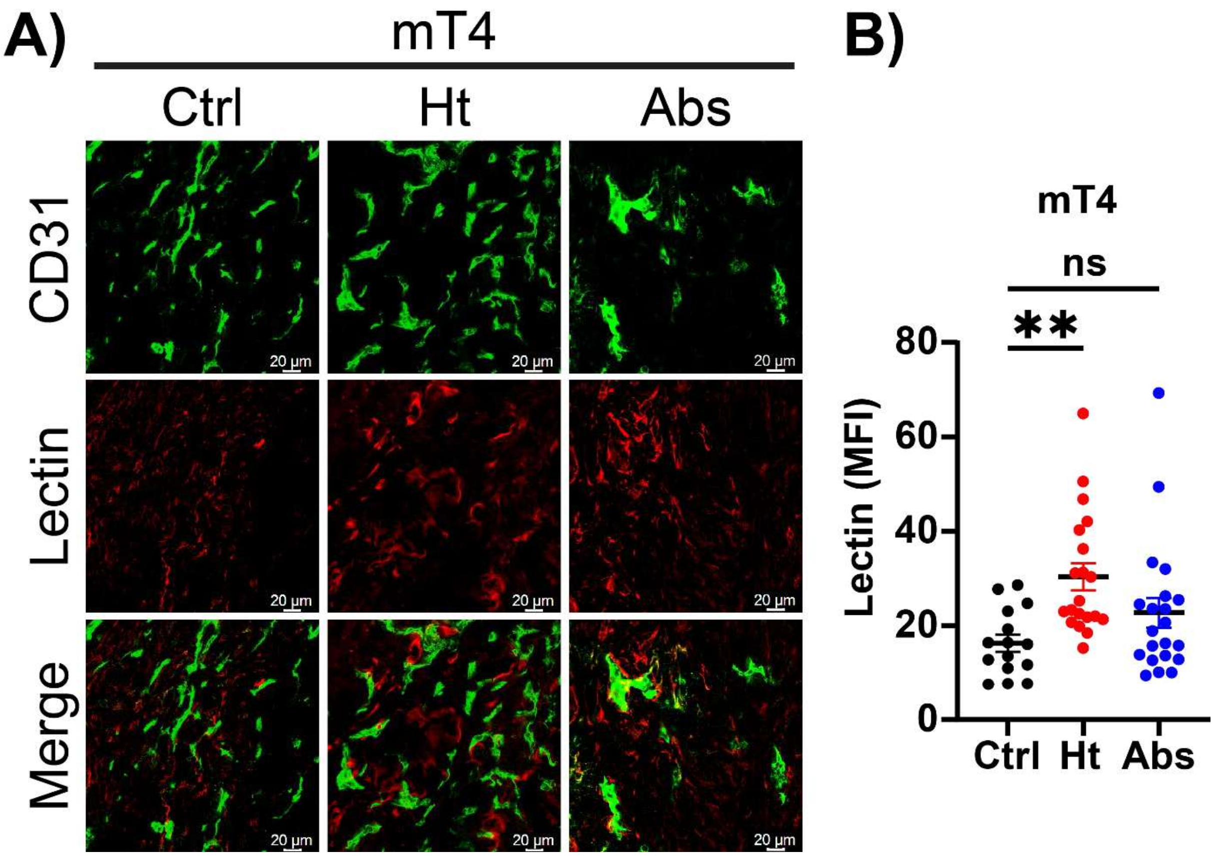


**Supplemental Figure S4. Histotripsy increases tumor perfusion.** (**A**) Co-staining of CD31 and lectin in control and treated mT4 tumors. (**B**) Quantification of lectin from (**A**) using MFI (ns = not significant; **p < 0.01; ****p < 0.0001; error bars represent standard error of the mean (SEM)).

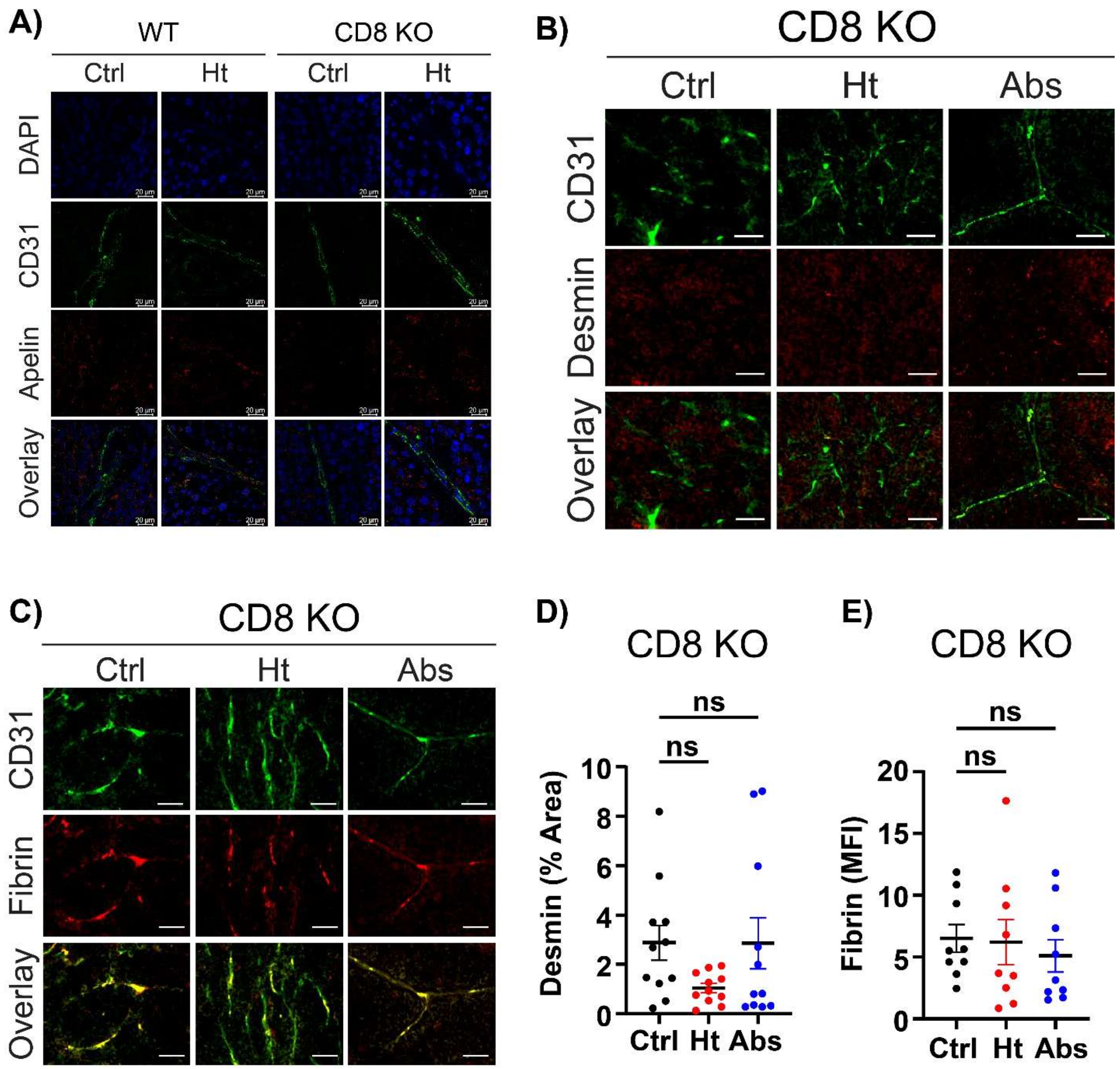


**Supplemental Figure S5. CD8+ T cells are necessary for histotripsy-induced vascular remodeling.** (**A**) IF staining of tumor endothelium (CD31) and pericyte (Apelin) of wild-type (WT) and CD8-knockout (KO) mice that were treated with sham or partial histotripsy. (**B–C**) Co-IF staining of CD8 KO mice with CD31 and Desmin and CD31 and Fibrin to indicate vessel maturity and leakiness, respectively. Scale bars represent 50μm. (**D**) Quantification of Desmin+ endothelium from (**B**). (**E**) Quantification of extravascular fibrin from (**C**) (ns = not significant; *p < 0.05; error bars represent standard error of the mean (SEM)).

**Table 1**. *Fold change of total vessel density (CD31+ area per mm$^2$) in different tumor-types.*

| **Tumor-type** | **Total vessel density (fold decrease)** | | |
|---|---|---|---|
| | Ctrl | Ht | Abs |
| **B16F10** | 1.0 | 2.7**** | 3.1**** |
| **Hepa1-6** | 1.0 | 7.4**** | 2.9** |
| **mT4** | 1.0 | 2.5*** | 5.0**** |
| *P values show comparison with the control tumors; **p < 0.005; ***p < 0.001; ****p < 0.0001 (One-way ANOVA with Dunnett's multiple comparisons test).* | | | |

**Table 2**. *Fold change of Apelin+ tumor endothelium in histotripsy-treated tumors.*

| **Tumor-type** | **Apelin coverage (fold increase)** | | |
|---|---|---|---|
| | Ctrl | Ht | Abs |
| **B16F10** | 1.0 | 2.4** | 1.7 |
| **Hepa1-6** | 1.0 | 3.8**** | 3.3**** |
| *P values show comparison with the control tumors; **p < 0.005; ****p < 0.0001 (One-way ANOVA with Dunnett's multiple comparisons test).* | | | |

**Table 3**. *Fold changes of Desmin and Fibrin expressions in different tumor-types after histotripsy.*

| **Tumor-type** | **Desmin expression (fold increase)** | | | **Fibrin expression (fold decrease)** | | |
|---|---|---|---|---|---|---|
| | Ctrl | Ht | Abs | Ctrl | Ht | Abs |
| **B16F10** | 1.0 | 3.9** | 3.1* | 1.0 | 1.1 | 1.9 |
| **Hepa1-6** | 1.0 | 5.4*** | 2.3 | 1.0 | 2.7** | 5.1*** |
| **mT4** | 1.0 | 2.2* | 0.6 | 1.0 | 5.7*** | 4.1*** |
| *P values show comparison with the control tumors; *p < 0.05; **p < 0.005; ***p < 0.001; ****p < 0.0001 (One-way ANOVA with Dunnett's multiple comparisons test).* | | | | | | |

**Table 4**. *Summary of vascular remodeling in different tumor-types following histotripsy.*

| **Tumor-type** | | **Reduction of vessel density** | **Vascular pruning** | **Vascular integrity** |
|---|---|---|---|---|
| **B16F10** | Treated | + | + | - |
| | Off-target | + | + | - |
| **Hepa1-6** | Treated | + | + | + |
| | Off-target | + | + | + |
| **mT4** | Treated | + | - | + |
| | Off-target | + | - | + |

**Table 5**. *Fold changes of intratumoral cell density and blood vessel diameter in histotripsy-treated tumors.*

| Tumor-type | Tumor cell density (fold decrease) | | | Blood vessel diameter (fold increase) | | |
|---|---|---|---|---|---|---|
| | Ctrl | Ht | Abs | Ctrl | Ht | Abs |
| **B16F10** | 1.0 | 1.3**** | 1.3**** | 1.0 | 2.0**** | 1.6**** |
| **Hepa1-6** | 1.0 | 1.5**** | 1.1 | 1.0 | 1.5**** | 1.1 |
| **mT4** | 1.0 | 1.5**** | 1.1 | 1.0 | 1.7**** | 1.5**** |
| **McA-RH7777** | 1.0 | 1.7**** | N/A | 1.0 | 1.6**** | N/A |
| *P values show comparison with the control tumors; ****p < 0.0001 (One-way ANOVA with Dunnett's multiple comparisons test; McA-RH7777 was analyzed using unpaired t-test).* | | | | | | |

**Table 6**. *Fold change of endothelial CXCR4 MFI in different tumor-types.*

| Tumor-type | Endothelial CXCR4 MFI (fold decrease) | | |
|---|---|---|---|
| | Ctrl | Ht | Abs |
| **B16F10** | 1.0 | 4.1* | 2.0 |
| **Hepa1-6** | 1.0 | 1.3 | 1.1 |
| **mT4** | 1.0 | 4.6** | 5.3*** |
| *P values show comparison with the control tumors; *p < 0.05; **p < 0.005; ***p < 0.001 (Two-way ANOVA with Tukey's multiple comparisons test).* | | | |

**Table 7**. *Fold change of the Angiopoietin1/2 ratio in histotripsy-treated tumors.*

| Tumor-type | Ratio of Angpt1/2 (fold increase) | | |
|---|---|---|---|
| | Ctrl | Ht | Abs |
| **B16F10** | 1.0 | 2.2* | 1.0 |
| **Hepa1-6** | 1.0 | 2.7** | 1.2 |
| **mT4** | 1.0 | 1.8 | 1.5 |
| *P values show comparison with the control tumors; *p < 0.05; **p < 0.005 (One-way ANOVA with Dunnett's multiple comparisons test).* | | | |
